# Proteome-level assessment of origin, prevalence and function of Leucine-Aspartic Acid (LD) motifs

**DOI:** 10.1101/278903

**Authors:** Tanvir Alam, Meshari Alazmi, Rayan Naser, Franceline Huser, Afaque A. Momin, Katarzyna W. Walkiewicz, Christian G. Canlas, Raphaël G. Huser, Amal J. Ali, Jasmeen Merzaban, Vladimir B. Bajic, Xin Gao, Stefan T. Arold

## Abstract

Short Linear Motifs (SLiMs) contribute to almost every cellular function by connecting appropriate protein partners. Accurate prediction of SLiMs is difficult due to their shortness and sequence degeneracy. Leucine-aspartic acid (LD) motifs are SLiMs that link paxillin family proteins to factors controlling (cancer) cell adhesion, motility and survival. The existence and importance of LD motifs beyond the paxillin family is poorly understood. To enable a proteome-wide assessment of these motifs, we developed an active-learning based framework that iteratively integrates computational predictions with experimental validation. Our analysis of the human proteome identified a dozen proteins that contain LD motifs, all being involved in cell adhesion and migration, and revealed a new type of inverse LD motif consensus. Our evolutionary analysis suggested that LD motif signalling originated in the common unicellular ancestor of opisthokonts and amoebozoa by co-opting nuclear export sequences. Inter-species comparison revealed a conserved LD signalling core, and reveals the emergence of species-specific adaptive connections, while maintaining a strong functional focus of the LD motif interactome. Collectively, our data elucidate the mechanisms underlying the origin and adaptation of an ancestral SLiM.

## INTRODUCTION

To a large extent, cellular signal transduction networks rely on the recognition of short linear motifs (SLiMs) by their cognate ligand binding domains^1^. These motifs are contained on a single contiguous amino acid stretch of typically less than 15 residues, and do not require to be embedded in a three-dimensional protein framework to be functional. The binding energy of many SLiMs is dominated by only a few residues, resulting in moderate to low binding affinities (with dissociation constants *K*_*d*_’s of 1–150 μM) that are ideal for mediating transient signalling interactions ^2^. On the one hand, this characteristic facilitates emergence and diversification of SLiMs, and hence the restructuration and evolutionary adaptation of an organism’s interactome ^3^. On the other hand, the resulting sequence motif degeneration and binding promiscuity hamper our capacity to computationally identify SLiMs and their biologically relevant binding partners ^4,5^.

LD motifs were first described in 1996 by Brown et al. as novel SLiMs that associate paxillin with the cell adhesion proteins vinculin and the focal adhesion kinase (FAK) ^6,7^. Paxillin and its family members Hic-5 (also called TGFB1I1 or ARA55) and leupaxin ^8-10^ contain in their N-terminal region four or five of these motifs, named after the first two amino acids of their sequence consensus LDXLLXXL ^7^ (Fig. 1A,B). The N-terminal LD motifs, together with other protein-protein interaction sites located in this region, orchestrate the dynamic assembly of different signalling complexes^9^. The C-terminal region of paxillin family proteins, which contains four double zinc-finger lin-11, isl-1, mec-3 (LIM) domains, mediates recruitment to integrin clusters at sites of cellular adhesion. The LIM domains also mediate nuclear localisation and nuclear receptor interactions (reviewed in ^10^). Using these diverse protein interaction motifs, paxillin family proteins establish a communication between cell adhesions and the nucleus, functionally linking gene expression with cell attachment ^11-13^. Thus, paxillin family members play important roles in embryonic development, epithelial morphogenesis and the immune response ^14^. As mediators of cellular motility and survival, paxillin family proteins are also key factors governing associated pathological conditions, such as inflammation, cardiovascular disease, and the development, spread and metastasis of tumours ^9,10,12,14-16^. Moreover, paxillin LD motifs are targets of a subset of human papilloma viruses, which cause cervical cancers^17,18^.

**Figure 1:**
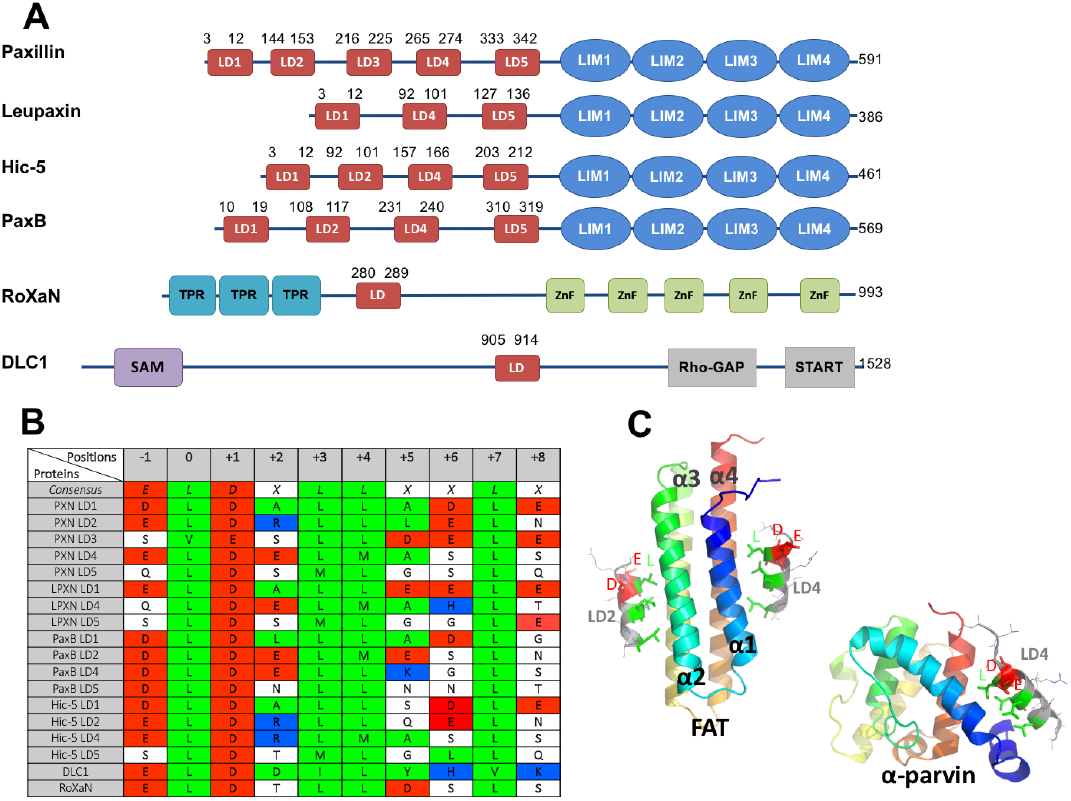
Overview of LD motifs. (A) Schematic representation of human paxillin family members (paxillin, leupaxin and Hic-5); PaxB is an orthologue from *Dictyostelium discoideum*. RoXaN and DLC1 both contain one LD motif. In addition, RoXaN has TPR and zinc finger (ZnF) domains. DLC1 has a SAM (sterile α-motif) domain, a GAP domain for small Rho family GTPases, a lipid-binding STAR-related lipid transfer domain. (B) Sequence alignment of selected known LD motifs. Sequence positions are numbered with respect to the first leucine of the LD motif (numbered 0). Acidic (red), basic (blue) and hydrophobic (green) residues are highlighted. PXN: paxillin. LPXN: leupaxin. All sequences are from human proteins, except for DLC1 (mouse) and PaxB (*D. discoideum*). (cC) Structure of LD motifs bound to FAK FAT and α-parvin. Ribbon diagrams of FAT and α-parvin are color-ramped from blue (N-terminus) to red (C-terminus). LD motifs are shown in gray, with key residues shown as stick models in wither green (hydrophobic) or red (acidic). Position −1 (D), 0 (L) and+1 (E) are labelled.

More than a dozen proteins were shown to interact with paxillin family LD motifs, using LD motif binding domains (LDBDs) of at least six different domain architectures (reviewed in^10^). Most of these proteins are important players in membrane-proximal intracellular structures that connect cell-surface receptors with signalling and/or the cytoskeleton, mostly in focal adhesions or similar structures (FAK, PYK2, vinculin, talin, GIT, parvin, Bcl-2)^6,16,19-24^. Other proteins are involved in nuclear export of proteins (XPO1, also called CRM1) or the transport of mRNA (PABP1) for mRNA delivery to sites of cellular adhesion^11,25,26^. In all cases where experimental 3D structures are available, LD motifs form amphipathic helices that use the hydrophobic side chains (<u>L</u>DX<u>LL</u>XX<u>L</u>) of the motif to dock onto an elongated hydrophobic patch on LDBDs, while the negative charge (L<u>D</u>XLLXXL) forms ionic interactions with basic charges next to the hydrophobic patch (Fig. 1C) ^10,19,22,24^.

The biological importance of LD motifs, and the discovery of more than a dozen LD motif– interacting proteins (many of which have evolved different LDBD folds) ^10^, motivated efforts to discover LD motifs outside the paxillin family. However, to date, motifs of the LDXLLXXL consensus were only experimentally confirmed in two cellular proteins: the deleted in liver cancer 1 (DLC1) tumour suppressor gene^24,27^, and the rotavirus ‘X’-associated non-structural protein (RoXaN)^25^. In addition to these, gelsolin (an actin binding, severing, and capping protein mediating osteoclastic actin cytoskeletal organisation) was found to bind the LDBD of PYK2 through a similar motif (LDXALXXL) ^28^. Some authors further include motifs from the E6 associated protein (E6AP) and the E6 binding proteins (E6BP), two proteins that interact with E6 proteins of a subset of papillomaviruses. However, these motifs deviate from the LD motif consensus sequence (E6AP: LQELLGEE; E6BP: LEEFLGDYR) and/or structure (E6BP; discussed below) ^17,18,29-31^.

The fastest and most comprehensive way to search for LD motifs on a proteome-wide level would be through bioinformatic approaches. However, current algorithms produce too many false positives to be useful. We, therefore, developed an algorithm specific for LD motifs, which faced two challenges. Firstly, identification of LD motifs requires the (normally unknown) 3D structural context of candidate sequences. Secondly, the number of *bona fide* LD motifs is too low for classical machine-learning approaches. We overcame these challenges through the development of a support vector machine (SVM)-based model to estimate the structural content of a candidate sequence and the use of experimental feedback. We show that the resulting <u>LD m</u>otif finder (*LDMF*) tool enables a proteome-scale prediction of LD motifs without a high rate of false positives, while still being able to distinguish LD motifs from highly similar short helical interaction motifs. Combined with experimental validation we used *LDMF* to assess the prevalence and function of LD motifs in the human proteome, to investigate the motif’s origin in unicellular eukaryotes, and probe its use in bacterial and viral pathogens. Our integrative method can serve as a guide for the identification of other low-abundance SLiMs.

## RESULTS

### Features that characterise *bona fide* LD motifs *in silico.*

To determine features that characterise LD motifs *in silico*, we analysed known LD motif–containing proteins using algorithms to predict protein disorder (MetaPrDos^32^), as well as secondary (PSIPRED^33^) and tertiary protein structures (SwissModel^34^ and RaptorX^35^). Paxillin family LD motifs were located in regions predicted as disordered (**Supplementary Fig. 1**). Of the five paxillin LD motifs, only LD2 was predicted as ‘not disordered’, in agreement with previous NMR analyses showing that LD2 forms a stable α-helix in solution, whereas LD4 does not^36^. The LD motifs of RoXaN and DLC1 were also predicted to be ‘ordered’ regions within a largely disordered protein segment. Secondary structure prediction assigned a significant α-helix likelihood to all paxillin LD motifs, in agreement with structural studies, showing that DLC1 and paxillin LD motifs 1, 2 and 4 adopt helical conformations upon ligand binding (Fig. 1C) ^10,19,22,24^. Gelsolin’s C-terminal LD-like motif is predicted to be α-helical, preceded by a coiled region, in agreement with the crystal structure of the assembled form of gelsolin^37^ (**Supplementary Fig. 1).** Of the atypical LD-motifs, E6AP is predicted to form an α-helix within an unstructured region (no 3D model can be established), whereas the E6BP LD motif is part of a calcium-binding EF-hand (**Supplementary Fig. 2**), in agreement with experimental data^38^. The E6BP LD motif only binds to the viral E6, not to other cellular LDBDs, and the structural basis of this interaction remains to be determined. *Bona fide* LD motifs are therefore computationally identifiable as short α-helical segments within disordered protein regions.

### Assessment of the representative LD motif set

In 1998, Brown and colleagues used the degenerate sequence pattern (L,V)(D,E)X(L,M)(L,M)XXL to search sequence databases for LD motifs. They found this sequence pattern in a diverse array of proteins, suggesting that LD motifs are relatively abundant^7^, and presented a representative set of 18 LD motif–containing proteins. Based on data and algorithms published since then, we critically assessed these predictions using disorder and structure prediction algorithms (**Supplementary Fig. 2**). 16 out of the 18 suggested LD motifs were predicted to be an integral part of a folded protein domain. In 15 out of these 16 cases, the hydrophobic patch of the suggested LD motif is inaccessible to solvent and hence ligands. In the one remaining case (LTK), the suggested LD motif is part of the catalytically important αC helix of a protein kinase domain. Thus, unless unlikely large unfolding events occur, these 16 putative motifs cannot function as LD motifs despite containing the correct sequence pattern. For the remaining two of the 18 proteins, the suggested LD motif sequence is located in a flexible region. However, in one case (Eph-2) the putative LD motif is part of a signalling peptide that is cleaved *in vivo*, and hence an unlikely candidate. The remaining LD sequence from chicken tensin was a plausible candidate, being located in an unstructured region and implicated in focal adhesions ^39^. But the synthetic tensin LD motif peptide failed to show binding to two broad-specificity LDBDs (the FAK FAT domain and the second CH domain of α-parvin) using isothermal titration calorimetry (ITC), microscale thermophoresis (MST) and fluorescence anisotropy (FA). We additionally tested the suggested LD motif sequences of the E3 ubiquitin-protein ligase hyd for which our predictions were least compelling, but did not observe binding to FAT (**Supplementary Fig. 3**). Given the high number of false positives of the pattern search (which currently retrieves >6000 sequences in the human proteome), we tested ANCHOR^40^ in combination with the LD motif pattern from the Eukaryotic Linear Motif (ELM) database^41^. ANCHOR predicts interaction motifs that reside in disordered regions and are likely to gain order upon binding, which is a characteristic of LD motifs. ANCHOR suggested 417 LD motifs in the human proteome but also classified the 18 Brown et al. LD motifs as true LD motifs, although our combined computational and experimental analysis provided strong evidence that this is not the case.

### Development of a computational LD motif identification method

Given that LD motif prediction was an unsolved problem, we decided to build a specific tool (see Methods). For this, we extended the 8-residue LD motif (LDXLLXXL) by one amino acid on each side into a 10-residue core motif (X^-1^L^0^DXLLXXLX^+9^; the first leucine is numbered as 0), because structural analysis shows that positions −1 and +9 can contact the LDBD surface, and hence may contain LD motif-specific information^31^. As a computable proxy for the (generally unknown) 3D structural context of candidate sequences, we used secondary structure predictions of the core sequence and of the 20 upstream and 20 downstream residues. Machine-learning further included the amino acid sequence for the 10-residue core and 20-residue flanking sequences and the Amino Acid Index (AAindex) to extract volume, hydrophobicity and electric charge for the 10-residue core.

An additional difficulty for machine-learning was the imbalance and definition of the positive and negative datasets. The initial positive set contained only the 18 experimentally confirmed LD motifs from six proteins: four or five from each paxillin family protein (human paxillin, leupaxin and Hic-5 and dictyostelium paxillin-B); one from DLC1 and one from RoXaN (Fig. 1A). The negative dataset (i.e. all UNIPROT 10-mer sequences that are not LD motifs) is million-times larger than the positive set, yet undefined because the occurrence of LD motifs is unknown. We first extracted the known LD motifs from the sequences of the six LD motif proteins as the positive set and used the remaining sequence fragments as the negative set (Fig. 2). Sequence, secondary structure and AAindex features of these sets were used to build an initial model (M1). We then applied M1 to identify putative LD motifs in close homologues of our six positive-set proteins, resulting in additional 40 LD motif sequences that we manually checked and added to the positive set. The negative dataset was enhanced by using 2754 sequences from high-resolution structures deposited at the PDB (none of which contained LD motifs) to capture instances where the LD motif sequence is part of a folded domain (and hence not a functional LD motif). These training sets were used to build another model (M2), with which we scanned the human proteome (20159 sequences). All predicted novel LD motifs were synthesized as peptides and used in *in vitro* binding experiments (described below). The sequences that showed clear binding were added to the positive set, and input into the second phase of the model building. The resulting final *LDMF* tool identifies LD motifs with high sensitivity (88.88%) and accuracy (99.97%; **Supplementary Table 1**). In our evaluation, *LDMF* rejected all 18 motifs previously suggested by Brown et al. ^7^, in agreement with our computational and experimental assessment.

**Figure 2:**
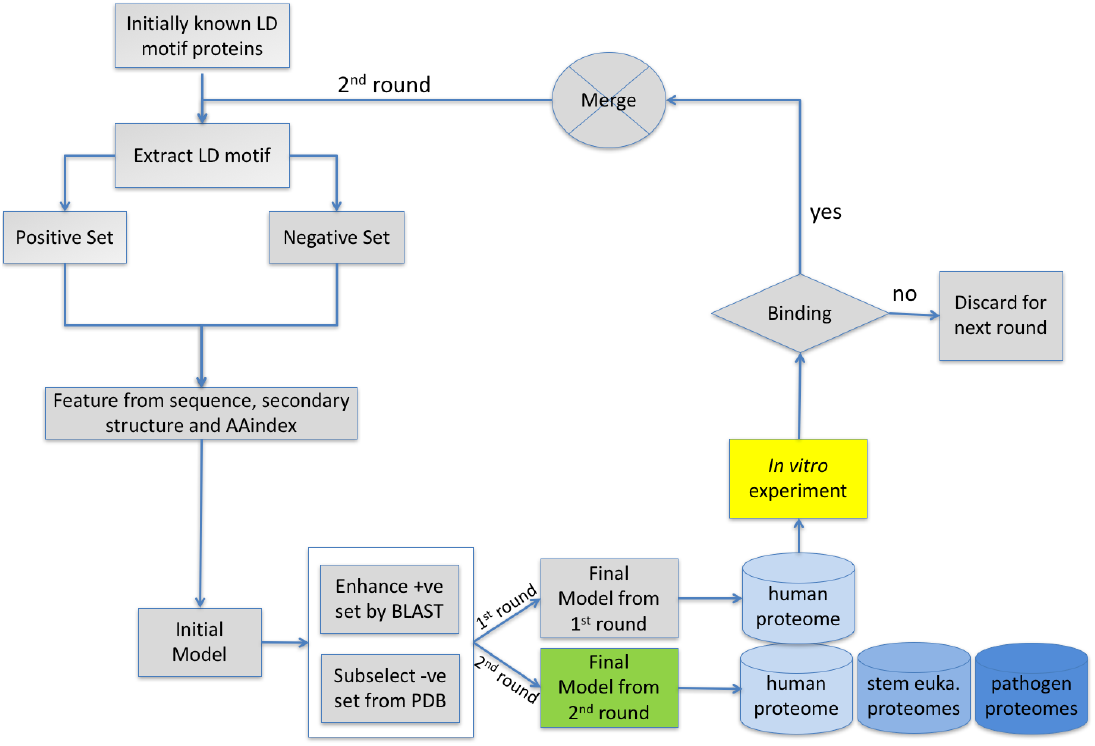
Flowchart of the set of operations performed for building the *LDMF* tool. The LD motif regions of the initially known LD motif proteins were extracted and used as initial positive data set. The remaining regions of the known LD motif proteins were considered as negative set. For both data sets, we used the sequence, secondary structure and amino acid physiochemical properties to represent those areas as a feature vector in our computational model. Using these features, we build our initial model. This initial model was used to enhance the positive (+ve) and negative (-ve) dataset. After integrating the new dataset with the initial dataset, the final model for round 1 was built. All predictions were tested experimentally and sequences that displayed LDBD binding were used as known LD motifs in a second round. The final model of the second round was used to scan proteomes from human, stem eukaryotes and a selection of bacterial and viral pathogens.

**Table 1:** Summary of experimental binding assays between putative LD motifs and selected LDBDs. The LD motifs are coloured according to: positive controls (green), negative controls (red), highly-likely (blue), less likely (orange), least likely (yellow) and the motifs discarded in round 1 (grey). Values indicate the *K*_*d*_ in μM for direct anisotropy (DA), microscale thermophoresis (MST) and isothermal calorimetry (ITC). ‘N’ indicates that no confident *K*_*d*_ could be derived from fitting the data. ‘-’ signifies that this measure was not performed. For anisotropy competition assay (ACA), and differential scanning fluorimetry (DSF) where no *K*_*d*_ could be determined, results are given as relative difference, or indicate *Tm* shifts, respectively. For ACA and DSF values, a t.test of significance was performed (n=3), were the null hypothesis is rejected with 95% (*), 99% (**) and 99.9% (***) of confidence. For *K*_*d*_ values, errors are indicated as SEM.

| | FAT | | | | | α-parvin | | | | GIT1 | | | |
| --- | --- | --- | --- | --- | --- | --- | --- | --- | --- | --- | --- | --- | --- |
|  | ACA | DSF | DA | MST | ITC | ACA | DSF | DA | MST | DSF | DA | MST | ITC |
| LD4 | 0.22 ***<br>0.36 *** | 0.4 ***<br>1.1 *** | 7.1 ± 0.2 | 55.5 ± 5.9 | 19.7 ± 2.9 | 0.09 **<br>0.19 ** | -0.2<br>-0.2 | 24.5 ± 0.8 | 3.1 ± 0.2 | - | - | - | - |
| LD2 | - | 2.2 ***<br>3.1 *** | - | - | - | - | 0.3<br>1.2 *** | - | - | - | - | - | - |
| E41L5 | 0.02<br>0.12 *** | 1.6 ***<br>3.5 *** | 2.7 ± 0.2 | 13 ± 0.9 | 6 ± 1 | 0.14 ***<br>0.28 *** | 1 ***<br>1.8 *** | 28.2 ± 2.4 | 4 ± 0.4 | 0.6 ***<br>1 *** | 83.6 ± 3.6 | N | 39.3 ± 5.4 |
| LPP | 0.03 *<br>0.04 * | 0.3 ***<br>0.7 *** | 97.4 ± 16.4 | N | N | 0.04<br>0.08 * | 0.2 **<br>0.3 ** | N | 169 ± 19 | - | - | - | - |
| RGPA1 | 0.03<br>0.03 * | 0.01<br>-0.1 | 43.6 ± 2.2 | 42.5 ± 1.8 | N | 0.05 **<br>0.1 ** | 0.1<br>0.2 | 49.4 ± 3.9 | 25.7 ± 0.7 | -0.3 *<br>-0.2 | 88 ± 18 | 77.5 ± 4.9 | N |
| P2R3A | 0.14 **<br>0.23 *** | 0.7 ***<br>1.6 *** | 41 ± 6.6 | 90 ± 5 | 19.7 ± 4.2 | 0.15 ***<br>0.26 *** | 1 ***<br>1.7 *** | 23.3 ± 4.4 | N | - | - | - | - |
| CD158 | 0.03<br>0.03 | 0.5 ***<br>1.3 *** | 41.6 ± 5.2 | N | N | 0.17 ***<br>0.2 *** | 0.6 ***<br>0.9 *** | N | N | - | - | - | - |
| CP071 | 0.04 *<br>0.07 ** | 0.4 ***<br>1.2 *** | N | 17.2 ± 2.2 | N | 0.1 ***<br>0.2 *** | 0.7 ***<br>1.4 *** | 52 ± 6 | 32.3 ± 3.5 | - | - | - | - |
| NCOA2 | 0.05 **<br>0.08 *** | 0.3 **<br>0.7 * | N | N | - | 0.05 *<br>0.08 ** | 0.1 ***<br>0.2 ** | N | 251 ± 14 | -0.3<br>0.1 | N | - | - |
| NCOA3 | 0.08 ***<br>0.15 *** | 0.3 *<br>0.7 *** | N | 261 ± 22 | N | 0.11 ***<br>0.21 *** | 0.2 **<br>0.4 *** | N | N | -0.1<br>-0.1 | N | - | - |
| CAST | 0.02<br>0.01 | 0.2 **<br>0.4 ** | N | 178 ± 13 | - | 0.02 *<br>0.05 * | 0.03<br>-0.1 | N | 273 ± 12 | - | - | - | - |
| CREB3 | 0.09 **<br>0.13 *** | 0.1<br>0.7 *** | N | N | - | 0.08 ***<br>0.13 *** | 0.8 ***<br>1.3 *** | 101 ± 5 | 110 ± 7 | - | - | - | - |
| RGPA2 | 0.01<br>0 | 0.5 ***<br>1.2 *** | N | N | - | 0.06 **<br>0.1 ** | 0.2 **<br>0.2 | N | N | 0.1<br>0.4 ** | N | - | - |
| CH037 | 0.01<br>-0.01 | -0.2<br>-0.8 | N | N | N | 0.16 *<br>0.04 * | -0.2<br>-0.6 | N | N | - | - | - | - |
| FIP1 | 0.03 *<br>0.03 * | 0.1 *<br>0.6 *** | N | N | N | 0.04<br>0.02 * | 0.03<br>-0.1 | N | N | - | - | - | - |
| IBP2 | 0.05 *<br>0.03 * | 0.3 ***<br>0.5 * | N | N | - | 0.01<br>0.01 | 0.04<br>-0.03 | N | N | - | - | - | - |
| PCP2 | 0.05 **<br>0.03 | 0.1<br>0.3 ** | - | - | - | 0.04 *<br>0.06 *** | -0.01<br>0.04 | - | - | - | - | - | - |
| WHAMM | 0.04 *<br>0.04 | 0.1<br>0.4 *** | N | N | - | 0.04 **<br>0.05 ** | 0.1<br>0.2 * | N | N | - | - | - | - |
| Scramble | 0.03<br>0.03 | 0.1 **<br>0.2 ** | N | N | - | 0.03 *<br>0.03 * | -0.02<br>-0.1 | N | N | -0.1<br>-0.2 | N | N | - |

### Experimental testing of all predicted LD motif candidates

Experimental testing of our computational LD motif predictions was not trivial, because the binding affinities of peptide-mimics of LD motifs towards their biologically relevant ligands are very low, ranging from *K*_*d*_’s of a few micromols to more than hundred micromols (summarised in^10^). Such affinities are below the detection limit of many assays, and the high concentrations required may lead to sample aggregation or non-specific interactions (between fluorescent moieties, protein/peptides, and storage or measurement containers). The (in)direct multimerisation or clustering of LD motif proteins and/or LDBD proteins and the bivalency of some LDBDs (i.e. the FAT domains from FAK and PYK2) might potentiate interactions of full-length proteins in cells^10,16^. Yet, LD motif interactions are tightly controlled in the cell, and biologically relevant LD motif interactions may not be detected using pull-down assays with endogenous proteins^24^. Establishing the required cellular conditions for each potential ligand, however, was incompatible with proteome-wide approaches.

Therefore, we used several orthogonal *in vitro* binding methods to robustly assess the interaction between synthesised peptides and recombinant LDBDs. For initial high-throughput screening, we used three plate assays: 1) differential scanning fluorimetry (DSF) was chosen as a semi-quantitative label-free binding indicator; 2) a direct anisotropy (DA) assay with labelled candidate peptides was chosen to estimate the interaction affinity; and 3) an anisotropy competition assay (ACA) where unlabelled candidate peptides compete against fluorescently labelled known LD motifs, was chosen to assess whether the (unlabelled) candidate motifs bind to the same sites as the known LD motifs. For all candidates, we used microscale thermophoresis (MST) with labelled peptides as an orthogonal quantitative method. Isothermal titration calorimetry (ITC) was used as an additional label-free method in selected cases to provide an additional binding *K*_*d*_, or binding stoichiometry. Nuclear magnetic resonance (NMR) was used in special cases to map binding sites. Peptide sequences included four to eight flanking residues outside the 10-residue core sequence. These additional residues were chosen based on homology modelling, secondary structure and disorder predictions to include helix-capping residues and residues that might additionally contact the LDBDs. Peptides were synthesized with and without a FITC-Ahx N-terminal fluorescent label.

Different LDBDs bind known LD motifs with different levels of specificity^10^. Since *LDMF* has been trained on all known LD motifs, without any specific target LDBD selectivity, we tested the affinity of the candidate LD motifs towards two recombinant human LDBDs with broad binding characteristics: firstly, the four-helix FAT domain from FAK (residues 892-1052) that can bind two LD motifs simultaneously (on opposite sides of the helix bundle)^19,42^. With these two binding sites, FAT binds paxillin LD motifs 1, 2 and 4, but also DLC1 ^24,27^, and the non-LD motif CD4 endocytosis motif ^43^.Secondly, we used the second CH domain of α-parvin (α-parvin-CH_C_, residues 242-372), which is a helical domain structurally distinct from FAT domains. This CH domain has one LD motif binding site that interacts with paxillin LD1, 2 and 4 with *K*_*d*_ values of 50 – 140 μM ^22,23^.

Peptide mimics of paxillin LD4, which were used as positive controls, displayed micromolar *K*_*d*_ values for FAT and α-parvin as expected, and competed efficiently against labelled LD4 in ACA (Table 1, **Supplementary Fig. 4**). Although the presence of LD4 resulted in a significant change in melting temperature *Tm* in DSF with FAT, the *Tm* change with α-parvin was not significant compared to a negative control (a peptide with the scrambled LD4 sequence). This result led us to include an LD2 peptide as a positive control in DSF.

In our binding assays (Table 1, **Supplementary Fig. 4**), candidate LD motifs display medium-to-low affinities as observed for established LD motifs ^10,24^. The relatively large discrepancies between *K*_*d*_s derived from different assays in some cases, and the frequent incapacity of individual binding assays to detect and quantitate binding (‘N’ in **Table** 1) highlight the difficulties in measuring LD motif:ligand interactions and justify our use of up to five different experimental methods. These experimental features made it unreasonable to define a clear cut-off to separate binding from non-binding sequences. Therefore, we ranked the peptides into (1) highly likely (*K*_*d*_ values from 1-99 μM in quantitative methods, and significant signal in at least one of the qualitative methods); (2) less likely (*K*_*d*_ values 100 μM and significant signal in at least one of the qualitative methods); (3) least likely (all others). Sequences labelled as ‘least likely’ in our assays may however still bind to the LDBDs of other proteins.

### Scan of the human proteome using *LDMF*

In the first scan of the human proteome, the initial *LDMF* tool predicted 13 new LD motifs (**Supplementary Table 2**). Those with the strongest experimental affinity for FAT and α-parvin (EPB41L5, RALGAPA1, C16orf71, LPP) were included in the training set for the second round of model building. The final *LDMF* tool predicted eight LD motifs in addition to the training set, five that were already predicted in the first round (C8orf37, RALGAPA2, NCOA2, NCOA3, CAST), and three new ones (PPP2R3A, CCDC158and CREB3). The final *LDMF* effectively discarded the four first-round candidates with the lowest experimental binding signals (FIP1L1, WHAMM, IGFBP2, PCP2). Thus, *LDMF* suggested 12 new LD-motif containing proteins in the human proteome. *In silico*, these 12 LD-motif candidates showed the required features of *bona fide* LD motifs (α-helices located in unstructured regions, not in a folded core) **(Fig. 3A). Experimental testing supported six candidates as ‘highly likely’ (EPB41L5, PPP2R3A, RALGAPA1, C16orf71, LPP, CCDC158), four as ‘less likely’ (NCOA2, NCOA3,CAST, CREB3), and two as ‘least likely’ (C8orf37, RALGAPA2) (Table 1, Supplementary Fig. 4**). We additionally assessed the *LDMF* predictions using four computational methods, namely PrePPI (a Bayesian framework that combines structural, functional, evolutionary and expression information^44^),GeneFriends (an RNAseq-based gene co-expression network^45^), CoCiter (which evaluates the significance of literature co-citations^46^), and a GO term analysis. (**Supplementary Table 3**).

**Figure 3A:**
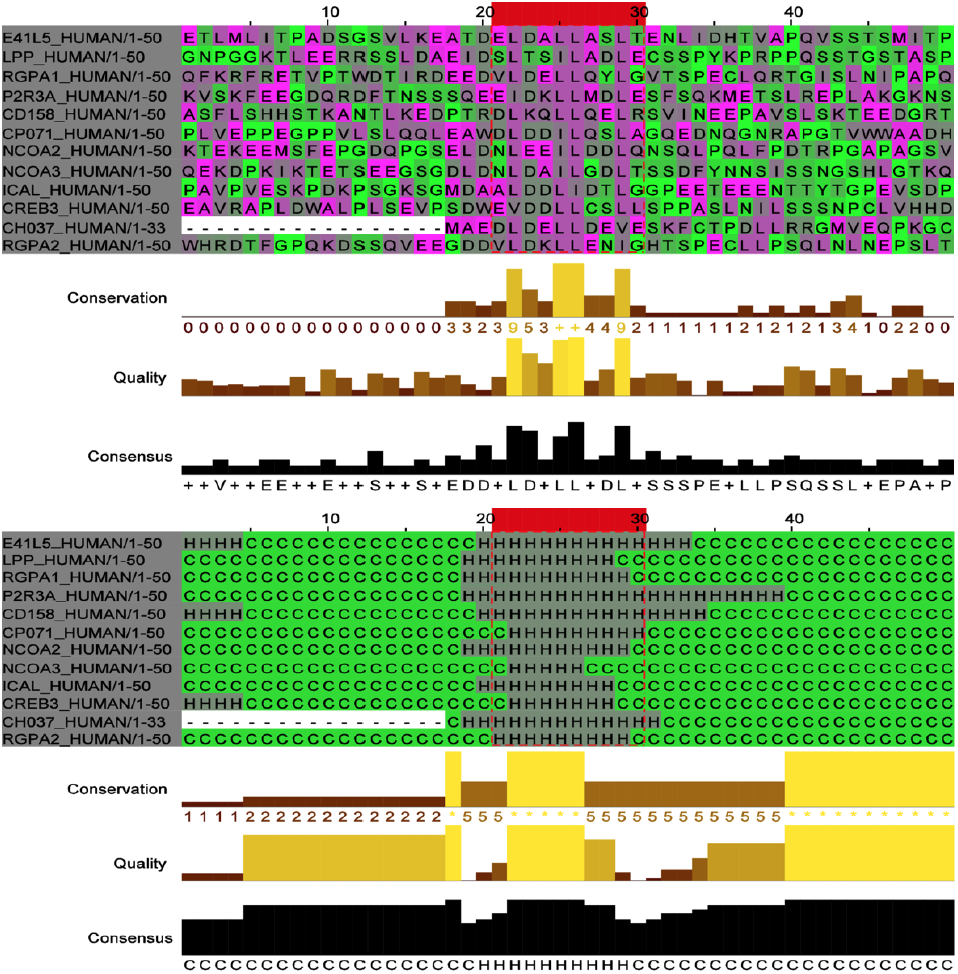
Sequence and secondary structure of predicted LD motifs in human proteins. The ten amino acids constituting the LD motif core are highlighted inside the red box. The twenty up-and down-stream residues of the flanking regions are shown. *Top:* amino acid sequences. *Bottom:* secondary structure. This figure was generated by Jalview.

**Figure 3B:**
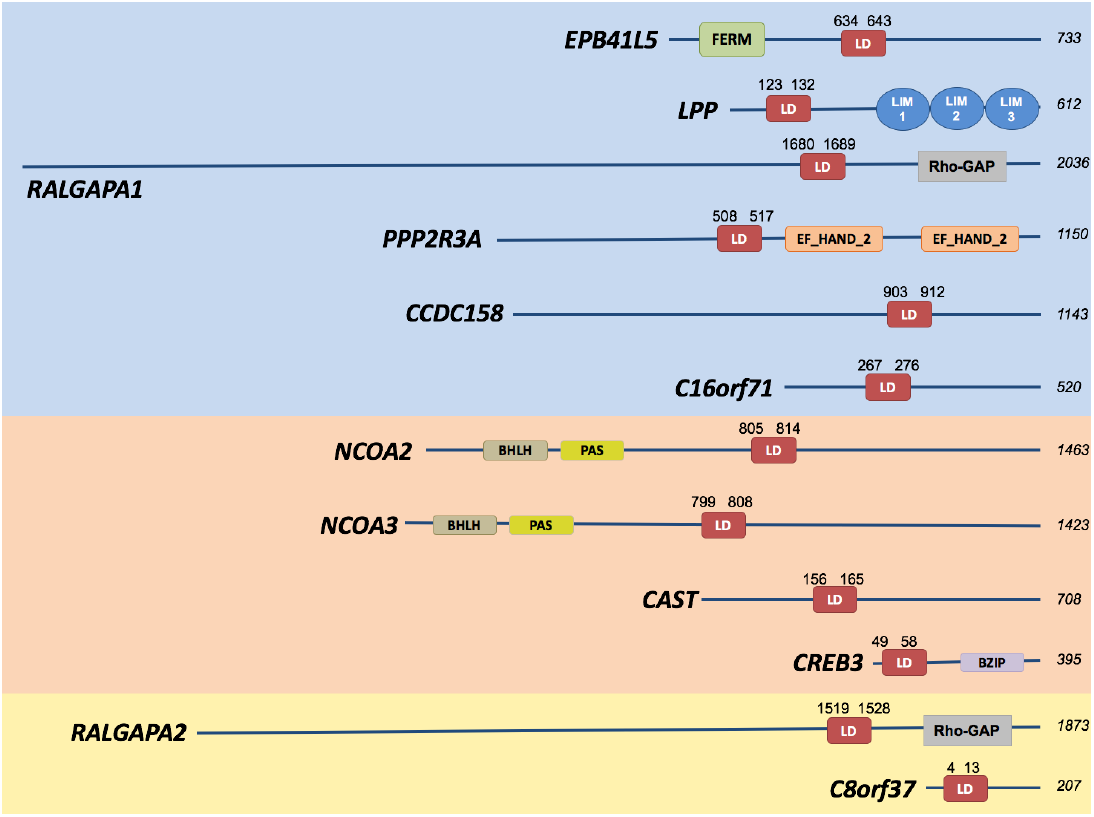
LD motif containing proteins identified in the human proteome. Protein length and positions of the LD motifs (residues −1 to + 8) are labelled. Additional domains are indicated by their PFAM name. Blue background: highly likely motifs; pale orange background: less likely motifs; yellow background: least likely motifs.

**Table 2:** Conservation of the non-paxillin LD motifs. Species investigated were chimpanzee, mouse, zebrafish, roundworm, fly, and yeast. ‘Distal’ refers to the evolutionary proximity to humans. *: with respect to the protein sequence in the most distal species. This table summarises results of **Supplementary Figure 7**.

| Gene name | Most distal protein homologue found in | residues identical /homologous to the human 10-aa LD motif * | LD motif is functional, according to LDMF (Y/N) * | Most distal functional LD motif to human (LDMF) |
| --- | --- | --- | --- | --- |
| *LDMF-identified* |  |  |  |  |
| EPB41L5 | shark | 8/9 | N | mouse |
| LPP | zebrafish | 10/10 | Y | zebrafish |
| RALGAPA1 | fly | 5/8 | Y | fly |
| PPP2R3A | shark | 7/10 | Y | shark |
| CCDC158 | shark | 3/5 | N | mouse |
| C16orf71 | mouse | 8/9 | Y | mouse |
| NCOA2 | zebrafish | 7/10 | Y | zebrafish |
| NCOA3 | zebrafish | 5/9 | N | mouse |
| CAST | shark | 6/6 | N | bird |
| CREB3 | bird | 6/9 | N | mouse |
| RALGAPA2 | fly | 5/6 | Y | fly |
| C8orf37 | shark | 9/10 | Y | shark |
| *Previously known* |  |  |  |  |
| RoXaN | shark | 6/8 | N | bird |
| DLC1 | shark | 7/8 | N | mouse |

### Proteins with highly likely LD motif sequences

#### Band 4.1-like protein 5 (EPB41L5)

The EPB41L5 peptide showed interactions in all methods used, and displayed highest affinities of all LD motifs tested (Table 1) towards both FAT and α-parvin. PrePPI probability scores for the interaction between EPB41L5 and FAK, PYK2 or Talin are >0.9 (**Supplementary Table 3**). EPB41L5 (also called YMO1 and LIMULUS, or yurt in Drosophila) contains an N-terminal FERM domain and a C-terminal flexible region, which harbours the predicted LD motif (residues 634-643) (Fig. 3B). EPB41L5/yurt is a critical regulator of the lateral membrane-associated cytoskeleton. It promotes FA formation by stimulating the interaction between paxillin and integrin.Although one report failed to observe an association between EPB41L5 and FAK or vinculin in TGF03B2;-treated NMuMG cells^47^, the high affinity and fitting cellular context of our study strongly support this LD motif.

#### Lipoma-preferred partner (LPP)

The LPP peptide showed qualitative and quantitative binding to both FAT and α-parvin. NMR chemical shift mapping supported a specific interaction with both binding sites on FAT (**Supplementary Fig. 5**). Interactions between LPP and vinculin, α-parvin, PYK2 or GIT1 are suggested by GeneFriends and CoCiter scores (**Supplementary Table 3**). LPP is a scaffolding protein that plays a structural role in the (dis)assembly of cell adhesions and may be involved in signal transductions from adhesion sites to the nucleus, thus affect activation of gene transcription^48,49^. LPP shows many similarities to paxillin family proteins: (i) Its N-terminal half is predicted to be an unstructured region harbouring proline-rich and phospho-tyrosine/threonine sequences in addition to the putative LD motif (residues 123-132); (ii) its C-terminal half contains three LIM domains. (iii) LPP shuffles between the nucleus and cytoplasm and is found at cell adhesions, including focal adhesions;(iv) the putative LPP LD motif sequence overlaps with a functional nuclear export signal (NES)^48-51^, supported by a PrePPI score of 0.98 for the LPP:XPO1 interaction. Moreover, LPP and FAK appear genetically linked ^50^. LPP and vinculin co-localise at focal adhesions and overexpression of the LPP LIM domains displaces LPP and vinculin from these structures ^48,51^.

#### Ral GTPase-activating protein subunit alpha-1 (RALGAPA1)

The RALGAPA1 peptide showed qualitative and quantitative binding to both FAT and α-parvin. PrePPI scores suggested RGPA1 interactions with FAK (0.76) and PYK2 (1.0). RALGAPA1 obtained a good co-expression score with the ARF GTPase–activating protein GIT2, and, in additional DA and MST experiments, bound with a *K*_*d*_ of 007E;80 μM to the GIT1 FAH domain (GIT1 and GIT2 are close homologues and have an identical LD motif binding site). RALGAPA1 (also called GARNL1 or TULIP1) is the catalytic α1 subunit of the heterodimeric RalGAP1 complex. RALGAPA1 functions as an activator of Ras-like small GTPases, including RalA and RalB ^52^. Activated RalA is involved in cell proliferation, migration, and metastasis. The suggested LD motif resides 100 amino acids upstream of the RapGAP domain, in a region predicted to be flexible (residues 1680-1689).

#### Serine/threonine-protein phosphatase 2A regulatory subunit B’ subunit alpha (PPP2R3A)

The PPP2R3A peptide showed qualitative and quantitative binding to both FAT and α-parvin. The predicted LD motif is located in a flexible region upstream of double EF hand domains (residues 508-517). PPP2R3A modulates substrate selectivity, catalytic activity and subcellular localisation of protein phosphatase 2A (PP2A)^53^. Indirect evidence suggests that PP2A promotes FAK phosphorylation^54,55^, interferes with the DLC1:FAK interaction ^56^, and is directly linked to focal adhesion proteins ^57^.

#### Coiled-coil domain-containing protein 158 (CCDC158)

The predicted LD motif in CCDC158 showed quantitative and qualitative binding to FAT and interacted specifically with the 1/4 site of FAT in NMR. We only measured significant qualitative binding to α-parvin. CCDC158 is an 1113-residue protein that contains 3 extended coiled-coil domains. The predicted LD motif region (residues 903-912) is located in a flexible region between the second and third coiled-coil. No literature is available for CCDC158.

#### C16orf71 (C16orf71)

C16orf71 is an uncharacterised protein of 520 residues. It is predicted to be mostly disordered, and the predicted LD motif region is located in the centre of the protein (residues 267-276) and showed qualitative and quantitative binding to both FAT and α-parvin.

### Proteins with less likely LD motif sequences

#### Nuclear receptor coactivator 2 (NCOA2)

NCOA2 is structurally mostly disordered and contains four nuclear receptor box (NR box) LXXLL motifs that mediate hormone-dependent co-activation of several nuclear receptors. A LLXXLXXXL motif in NCOA2 is involved in binding and transcriptional coactivation of CREBBP/CBP. The putative LD motif identified by *LDMF* (residues 805-814) is not part of these motifs, despite the similar consensus. Paxillin family proteins also bind to nuclear receptors, such as the androgen receptor and glucocorticoid receptor^10^. The region encompassing the putative NCOA2 LD motif is also predicted to function as an NES, akin to several paxillin LD motifs. NOCA2 obtained a strong CoCiter p-value (0.007) for association with PYK2, and has a large co-expression correlation with GIT2, but failed to show binding to GIT1 FAH in our additional experiments.

#### Nuclear receptor coactivator 3 (NCOA3)

Akin to NCOA2, NCOA3 is a scaffolding protein with many known interactors. NCOA3 has three NR box LXXLL motifs to bind to and co-activate several nuclear receptors, and a LLXXLXXXL motif to bind CREBBP/CBP. The predicted LD motif region (residues 799-808) is not part of already characterised motifs.

#### Calpastatin (CAST)

CAST is a specific inhibitor of the calcium-dependent cysteine protease calpain. The proposed CAST LD motif (residues 156-165) is a helical protein-protein interaction motif located in an otherwise disordered region; it uses its hydrophobic patch to bind to a helical subdomain of calpain, thus stabilising calpain in its inhibited form ^58^. Calpain regulates functions such as cell migration and anoikis (cell death following detachment), and among its substrates are Cas, talin, FAK and PYK2 ^59,60^. Calpain also associates with FAT ^60^. Hence a possible competitive interaction of the calpastatin LD motif with FAK and/or PYK2 interaction with CAST could potentially promote cleavage by calpain.

#### Cyclic AMP-responsive element-binding protein 3 (CREB3)

CREB3 is a single-pass transmembrane endoplasmic reticulum (ER)-bound transcription factor involved in the unfolded protein response, in cell proliferation and migration, tumor suppression and inflammatory gene expression. The predicted LD motif region (residues 49-58) is located in a flexible acidic transcription activation region downstream of a basic leucine-zipper (bZIP) domain (residues 174-237) and the transmembrane region (residues 255-271).

### Proteins with least likely LD motif sequences

#### Ral GTPase-activating protein subunit alpha-2 (RALGAPA2)

RALGAPA2 is the catalytic α2 subunit of the heterodimeric RalGAP2 complex. The putative LD motif (residues 1519-1528) lies in a poorly ordered region for which no 3D template is available. The RALGAPA2:PYK2 interaction has a PrePPI score of 0.93, shows medium-level co-expression with GIT2, but failed to show binding to GIT1 FAH.

#### C8orf37 (C8orf37)

C8orf37 is a poorly characterised protein of 207 residues, predicted to have a disordered N-terminal half and a C-terminal half with an α/03B2; fold. The putative LD motif (residues 4-13) is at the N-terminus. C8orf37 is widely expressed, with highest levels in brain and heart. C8orf37 is also expressed in the retina and mutations in this gene have been linked to an inherited retinal dystrophy.

### *LDMF*-identified LD motif proteins are functionally similar to *bona fide* LD motif proteins

Using gene ontology (GO) analysis we found that the distribution of GO semantic similarity between predicted and known LD motif proteins is significantly different from the distribution between predicted and all human proteins (p-value = 6.32e^-10^, Mann-Whitney test; **Supplementary Fig. 6**). Especially, GO terms with a dispensability value (i.e. the semantic similarity threshold at which the term was removed from the list and assigned to a cluster) of zero were the same for the predicted and known LD motif proteins for both categories, biological processes (BP: regulation of cell morphogenesis, biological adhesion and cell-substrate adhesion) and cellular components (CC: focal adhesion, basolateral plasma membrane and cell junction). These zero-dispensability terms were similar to those of the proteins containing LDBDs (BP: signal complex assembly, biological adhesion, and cell-substrate adhesion. CC: focal adhesion, basolateral membrane, cell junction and organelle). Our predicted LD motif proteins are therefore functionally highly similar to the known LD motif proteins and to LDBD-containing proteins.

### Identification of an inverse LD motif consensus

Surprisingly, the predicted and experimentally confirmed LD motif sequences for LPP and CCDC158 did not contain the consensus L^0^D motif. Rather, both sequences contained an LD/E motif in reverse orientation (D/E^+6^L). Titration with LPP or CCDC158 caused similar NMR chemical shift changes on 15N-labelled FAT titrated as LD2 or LD4, with LPP showing the preference for the site 2/3 (akin to LD4), whereas CCDC158 preferentially bound site 1/4 (akin to LD2) (**Supplementary Fig. 5**). These similarities strongly suggested that LPP and CCDC158 occupy the canonical binding sites, despite their reversed LD motif.

The pseudo-palindromic nature of the helical LXXLLXXL pattern would allow a reverse LD motif to engage similar electrostatic interactions to common LD motifs if the reverse motif bound in the opposite (-) orientation. Indeed, paxillin LD1 (DL^0^DXLLXDLE) binds CCM3-FAH and α/03B2;-parvin-CH_C_ in the opposite (-) orientation as LD2 and LD4 (+ orientation) ^19,22,23,61,62^, placing the D^+6^L^+7^E^+8^ motif in the position of the EL^0^D motif of LD2 and 4. We failed to produce co-crystals with sufficient diffraction power of these motifs bound to LDBDs. However, in support of the bioinformatics analysis, NMR data-guided *in silico* docking (HADDOCK) produced plausible low-energy reverse-orientation LPP:FAT and CCDC158:FAT models (Fig. 4).

**Figure 4:**
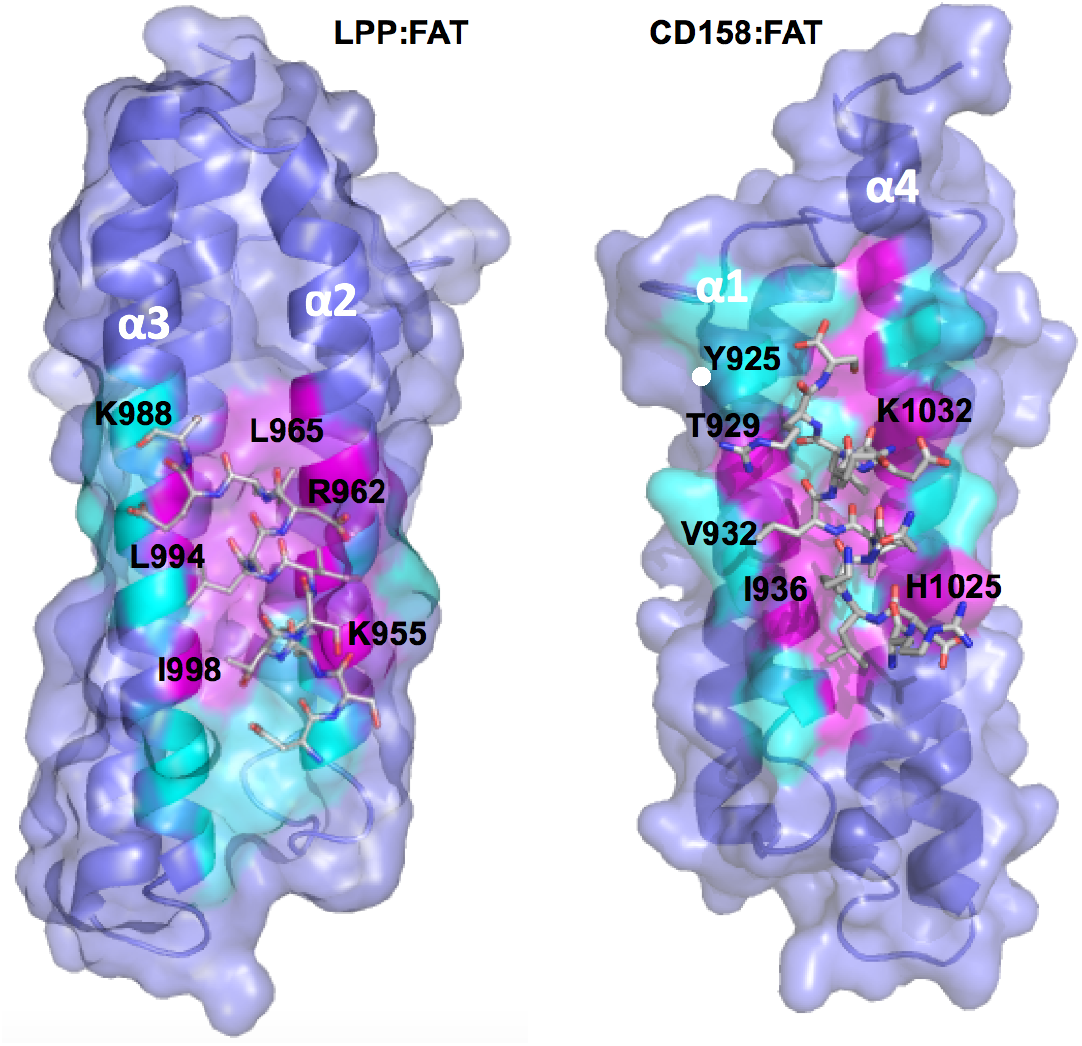
Suggested molecular basis for class II LD motif interactions with the FAK FAT domain. NMR-data guided docking of the LD motif peptides (stick models, with carbons colored in gray) derived from LPP (left) and CCDC158 (right) onto the FAK FAT domain (Secondary structures are shown in dark blue; molecular surface in light blue). Actively involved FAT residues are presented in magenta and passively involved residues are in cyan. Key residues on FAT for the interactions are labelled. The inverse consensus of these LD motifs is compatible with an opposite binding directionality compared to LD2 and LD4 motifs.

### Conservation of the non-paxillin LD motifs in animals

Paxillin homologues with functional LD motifs are found even in the amoeba *D. discoideum^63^*. We therefore used *LDMF* to evaluate the evolutionary conservation of the two known (RoXaN and DLC1) and twelve newly identified non-paxillin human LD motifs in chimpanzee, mouse, birds, fish, roundworm, insects, and yeast. None of these proteins had homologues in unicellular yeast, only two were found in flies (RALGAPA1, RALGAPA2) and most first appeared in fish (10) (Table 2, **Supplementary Fig. 7**). Where found, the *LDMF*-identified LD motif sequence generally was more conserved than the flanking residues, as expected for a SLiM within a disordered linker region. Seven out of the 14 proteins had functional LD motifs (according to *LDMF*) already in the first (most distal) species in which they were found. EPB41L5 showed a weak below-*LDMF* threshold LD motif signature in the shark, none in chicken, and a strong LD motif in mice. For the other genes, LD motifs appeared only in birds (2) or mammals (5). Given that some known LD motifs also function as NES, we additionally tested the degenerate LD motifs for NES function using the NetNES1.1 server ^64^. We found that in most species where the LD motif was not predicted to be functional (EPB4L5, CCDC15, NCOA3, CREB3, DLC1), this sequence was identified as functional NES. Exceptions were RoXaN and CAST, where the NES signal appeared above noise, but below threshold. In conclusion, the arsenal of non-paxillin LD motifs currently found in humans appears to have originated in multicellular animals, and mostly in vertebrates. In several instances, we observe evidence for the appearance of a functional LD motif *de novo* within a protein from a sequence that is expected to function as NES.

### Scan for LD motifs in stem eukaryotes using *LDMF*

Although paxillin and other LD-motif proteins regulate cell-matrix adhesion in metazoans, proteins of the integrin signalling pathway were discovered in the unicellular eukaryotic lineages ichtyosporea and filastera, supporting that LD motif signalling played already a role prior to the evolution of metazoans ^65^. We used *LDMF* to investigate the importance and evolution of LD motif signalling in the ten sequenced stem eukaryotes proteomes that were available, covering amoebozoa, apusozoa and unicellular ophistokonts.

Paxillin-like LD motif-containing proteins were identified in all but two species, *Monosiga brevicollis* and *Mortierella verticillata* (Fig. 5). These two species are of different ophistokont clades, each of which contains also species with paxillin homologues, suggesting that paxillin was selectively lost in *M. brevicolis* and *M. verticillata*. *LDMF* also identified between 1 and 20 non-paxillin LD motif proteins in each proteome. While some of the non-paxillin LD motif containing proteins were conserved between species, especially those proteins containing kinase domains, the majority was species-specific (39 out of 53 sequences; **Supplementary Fig. 8**).

**Figure 5:**
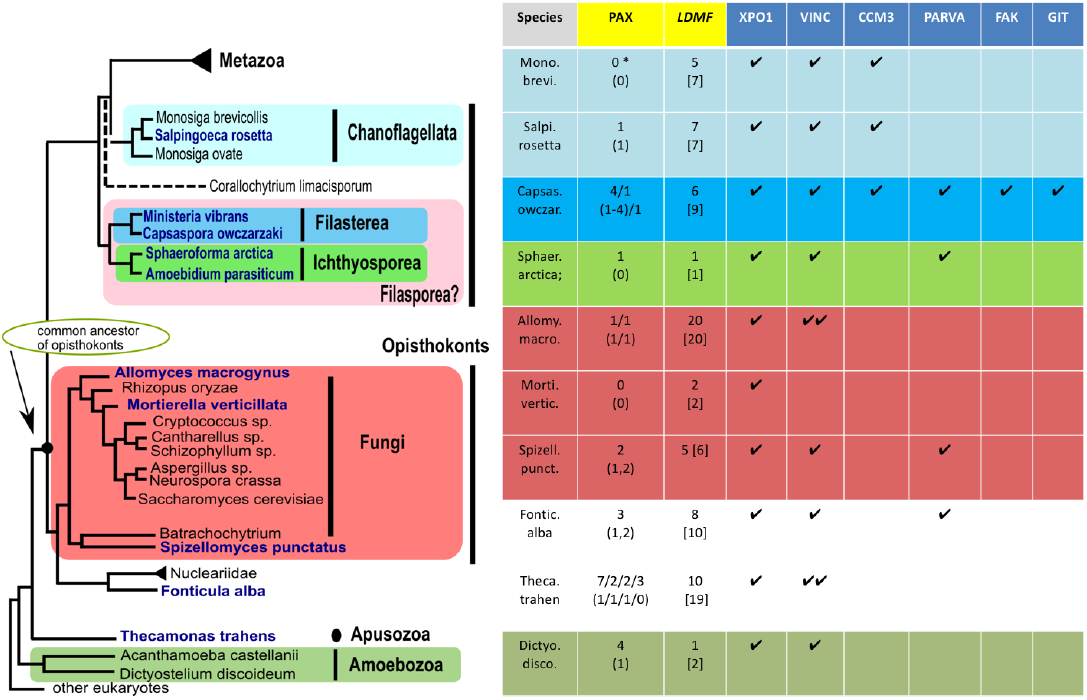
*LDMF*-predicted LD motifs and LDBDs in stem eukaryotes. *Left:* Evolutionary relation of the unicellular eukaryotes analysed. Figure adapted from the Broad Institute’s *Origin of Multicellularity* initiative. *Right:* The PAX column gives a summary of paxillin homologues, where the first row corresponds to the number of LD motifs found. In case there are several paxillin homologues in one species, the corresponding numbers are separated by ‘/’. An asterisk (*) indicates that this species contains a paxillin homologue but without LD motif. IN the second row, the numbers in brackets correspond to the positions of the LD motifs that are also predicted to be a functional NES. The *LDMF* column indicates in the first row the number of non-paxillin LD-motif containing proteins identified by *LDMF*, and in the second row in parenthesis the total number of LD motifs of a given species. The XPO1, VINC, CCM3, PARVA, FAK and GIT columns show the presence of genes homologous to exportin, vinculin, CCM3, *α*parvin, FAK and GIT, respectively. The number of ticks corresponds to the number of homologues found. The presence of a functional LDBD in these domains was assessed by sequence alignments and homology modelling.

We next used sequence homology searches and structural homology modelling to investigate the presence of functional LDBDs in these unicellular genomes. Of the six LDBD proteins investigated, only XPO1 and vinculin were present in all clades. XPO1 was found in all proteomes, as expected from its nuclear export function (Fig. 5; **Supplementary Fig. 8**). Vinculin was only absent in *M. verticillata*. The only two *LDMF*-identified non-paxillin LD motifs for this species were predicted to function also as NES (using the NetNES1.1 server), suggesting that *M. verticillata* had lost LD motif signalling and that the identified LD motifs were retained because of their NES function. Conversely, despite having lost paxillin, *M. brevicollis* encoded for two LDBD proteins (vinculin and CCM3) and at least five predicted LD motif proteins, suggesting that LD motif signalling remains used even in absence of paxillin.

Collectively these observations confirm the existence of LD motif-based signalling in unicellular unikonta. Our data suggest that this signalling pathway originated before the split between amoebozoa and ophistokonts, and emerged from a core module formed by paxillin and vinculin. Supported by our findings on the evolution of human non-paxillin LD motifs, we speculate that this pathway arose by co-opting NES, which can have an overlapping sequence motif. The relatively poor conservation of non-paxillin LD motifs in unicellular eukaryotes might thus be explained by interconversion of NES from non-related proteins during specification, and/or selective losses of LD motifs, resulting in the emergence of species-specific protein connectivities.

### LD motifs in pathogens

We further tested our model on several pathogens. We used proteins from bacteria (Mycobacterium tuberculosis, Salmonella typhimurium) and viruses (HIV, SIV, HCV). *LDMF* did not predict any LD motifs in proteins from these strains.

## DISCUSSION & CONCLUSION

### Proteome-wide insights in LD motif prevalence and function

SLiMs are core constituents of cellular signalling networks. Their proteome-wide detection and analysis across species can yield insights into the evolutionary mechanisms that allow creation and adaptation of a specie’s interactome. Being formed by a short and often degenerate consensus motif, proteome-wide detection of SLiMs is a challenging task. By combining computational and experimental approaches, we produced a bioinformatic tool, named *LDMF*, that allowed proteome-wide detection of LD motifs with high accuracy. With this tool, we identified twelve new LD motif-containing proteins in the human proteome. In comparison, only three new LD proteins were discovered using classical experimentation over the last 20 years. Our *LDMF* tool was capable of distinguishing between LD motifs and other similar motifs, such as the nuclear receptor box LXXLL motif.

Unexpectedly, our data suggested that an LD motif can also be formed in the inverse orientation, by L^+7^D/E^+6^ instead of L^0^D/E^+1^. Inverse LD motifs were found in LPP and CCDC158 that have L^0^T^+1^ and L^0^K^+1^ instead of L^0^D/E^+1^, and are likely to bind LDBDs in the opposite (-) orientation to common LD motifs. (+) and (-) binding poses also occur in interactions of proline-rich peptides ^66,67^ and NES ^68^. In analogy, we propose to name LDXLLXXL motif ‘class I’, and LXXLLXDL the ‘class II’ consensus. Given that paxillin LD1 (DL^0^DXLLXDLE) binds LDBDs in the (-) orientation, whereas leupaxin LD1 (ELDXLLXELE) binds in the same (+) orientation as paxillin LD2 and LD4 ^69^, the EL^0^D (or DL^+7^E) position appears to be the main determinant of the binding orientation in cases where acidic charges are flanking both L^0^ and L^+7^. Given the distribution of charges and lengths of side chains, paxillin LD3 (SLESLLDELE), and NCOA2 (NLEEILDDLQ) might also be class II motifs. *LDMF* has identified the class II consensus without specific training (however, LPP was included in the second-round training set), supporting that our tool generally is able to identify class II motifs.

Although we cannot exclude that several motifs remained undetected, the high sensitivity and experimentally verified accuracy of *LDMF* strongly support that the number of false negatives was low. Hence, less than 20 out of 007E;20,000 human proteins appear to have an LD motif. Humans have therefore less LD motif-containing proteins than previously thought, explaining the difficulties and slow pace in their detection. The number of LD motif proteins is similar to the number of currently known LDBD-containing proteins. Our GO analysis showed that LDBD proteins as well as previously known and *LDMF*-predicted LD motif proteins form a homogenous group in terms of biological processes (regulation of cell morphogenesis and adhesion) and subcellular cellular localization (basolateral plasma membrane, focal adhesions, and cell junctions).

Despite their function in regulating adhesions within multicellular organisms, previous analyses detected paxillin and other integrin-signalling components in the unicellular eukaryotic lineages ichtyosporea and filastera ^65^. Scanning the available proteomes of unicellular eukaryotes for LD motifs, we could confirm that LD motif signalling had already evolved before the split between opisthokonts, apusozoa and amoebozoa, more than 800 Myr ago. The conservation of homologues for paxillin, vinculin and XPO1 (exportin-1) across unicellular unikonta, and the observation that LD motifs can also function as NES would be in agreement with a hypothesis that LD motifs originated from NES motifs, and that paxillin and vinculin form the ancient core of the initial LD motif signalling pathway. A LD motif origin from NES is supported by the several cases where NES sequences within lower vertebrate homologues transformed into functional LD motifs in higher vertebrates. In many cases, the transformation into an LD motif (with or without losing the NES function) was achieved by only a few amino acid substitutions. Thus, conversion from NES into LD motifs (and possibly *vice versa*) appears to be an easy means for species-specific adaptation and rewiring of LD motif signalling in NES-containing but otherwise unrelated proteins. The possibility for achieving SLiMs with dual recognition by the nuclear export machinery and adhesion proteins may have provided opportunities for functionally linking cell attachment and motility with nuclear events, such as gene expression.

Our data illustrate how a conserved SLiM signalling core can become enriched with additional species-specific connections, while preserving a strong functional coherence. An analysis of the sequence conservation of well-known SLiMs that interact with SH2, SH3, PDZ and serine/threonine kinases had previously suggested that each of these SLiMs is particularly well conserved in proteins that are of the same functional group as the proteins containing the cognate interacting domains^3^. By demonstrating that a few point mutations may suffice to repurpose an existing protein sequence, our data support that evolution is much more likely to create or alter SLiMs, than to create or alter a three-dimensionally folded SLiM-interacting domain (which would require more mutations or domain duplication and specification). Thus, our data provide an illustration and rationale for how the functional focus of a particular SLiM is controlled by the functional range of its binding domain. In the case of LD motif signalling, the founding event might be the evolution of the vinculin tail domain’s capacity to interact with a subset of paxillin’s NES. Yet, many SLiMs are frequently recognized by more than one domain, for example SLiMs for SH2 or SH3 domains can also interact with PTB and WW domains, respectively ^67,70^. In LD motif signalling this phenomenon is particularly pronounced, where as many as six unrelated types of domain folds were found to bind to LD motifs (reviewed in^10^). Nonetheless, we found that LD motif proteins and LDBD-containing proteins span a functionally fairly coherent group. The evolutionary mechanisms that restrict the functional range of a SLiM in the case of multiple unrelated cognate domains appear incompletely understood.

Collectively, our integrated computational and experimental analysis shed light onto the origin, evolution and prevalence of a poorly understood SLiM that controls embryogenesis and immunity, and plays central roles in cancer cell spreading and cardiovascular diseases. By providing a means to investigate the LD motif interactome, our results might open new avenues for understanding and correcting pathological cell migration. Our *LDMF* tool is a convenient way to further interrogate the prevalence and evolution of LD motifs in newly sequenced species. More generally, the methodology used to produce our *LDMF* tool can be applied to other SLiMs for which only a few representatives are known. Given that *LDMF* is currently the only specific bioinformatic tool for predicting LD motifs, we made *LDMF* available as a web-based application at http://www.cbrc.kaust.edu.sa/ldmf/index.php.

## ACKNOWLEDGEMENTS

We acknowledge SOLEIL for provision of synchrotron radiation facilities for testing of hundreds of FAT:LD motif peptide crystals, and we would like to thank Martin Savko, Gavin Fox, William Shepard, Serena Sirigu, Leonard Chavas and Pierre Legrand for assistance in using beamlines PROXIMA I and PROXIMA IIA. We acknowledge support from the KAUST Imaging and Characterization Core Lab and the Bioscience Core Lab. We thank John Hanks and Craig Kapfer for their assistance with use of the Dragon Cluster, Robert Höhndorf for advice with the GO analysis, and Mariusz Jaremko and Lukasz Jaremko for advice with NMR data analysis. The research reported in this publication was supported by King Abdullah University of Science and Technology (KAUST) through the baseline fund and the Award No. URF/1/1976-04, URF/1/1976-06, URF/1/3007-01, URF/1/1976-02, BAS/1/1606-01-01 and#OSR-2015-CRG4-2602 from the Office of Sponsored Research (OSR).

## AUTHOR CONTRIBUTIONS

T.A., M.A, V.B.B, and X.G. designed and developed computational algorithms. V.B.B, X.B., J.M. and S.T.A designed and supervised research. T.A., M.A, V.B.B, X.G. and S.T.A. analysed computational data. R.N., F.H., K.W.W., C.G.C., M.A., A.A.M. and A.J.A. carried out experimental research and R.N., F.H., K.W.W., C.G.C., M.A., A.A.M., A.J.A. and S.T.A. analysed experimental data. R.H. contributed computational analysis of experimental data. T.A., M.A, V.B.B, X.G., R.N., A.A.M., F.H. and S.T.A wrote the manuscript. All authors critically read the manuscript.

## MATERIALS AND METHODS

### Overview on method development

To provide a high-accuracy algorithm for proteome-scale detection of LD motifs, we used a machine-learning approach that combines secondary structure prediction and physiochemical properties (from the AAindex database) of amino acids (AA) of the LD motif region. We formalise this problem as a binary-class classification problem of 10-mers, i.e., a subsequence of 10 amino acids, where LD motifs are considered as the positive set and 10-mers that are not LD-motifs are considered as the negative set. As the number of known LD motifs is small, it becomes an imbalanced dataset problem with a small positive and a huge negative dataset, which usually causes issues for classification methods.Therefore, we used a two-phase approach for building the prediction model. In the first phase, we considered the known LD motifs as the positive set and the remaining 10-mers extracted from these proteins as the negative set. As expected, these extracted 10-mers can be easily differentiated from the true LD motifs because they do not satisfy sequence patterns, secondary structure patterns or physicochemical patterns of the LD motifs. Therefore, a model trained based on such a trivial negative set may not be practically useful. Yet it provides us a rough predictor by assigning different weights to sequence-, secondary structure- and physiochemical-patterns. In a second phase, we used this predictor to obtain more difficult negative sets. This was done by selecting the 10-mers from the proteins in the Protein Data Bank (PDB) which satisfy some of these patterns according to the first predictor, but not all of them. We then used these new negative sets as well to train the final predictor. This results in an active learning framework to train an LD-motif predictor.

### Initial training data set

Our model uses information from protein sequence content of data-windows of length 10AA. Such windows are denoted as core windows. A core window is shifting one residue ahead. So, if a protein has a length L >= 10 AA residues, then there are L-10+1 possible candidate core windows to be considered by scanning the protein sequence as containing a putative LD motif.

By surveying the literature, the known LD motifs were found in Paxillin, Leupaxin, PaxB, Hic-5^8^, RoXaN ^71^, and DLC1^72^ and we selected these LD motifs. This resulted in a set of 18 genuine LD motif windows generated from six proteins. We denote this set as the set of known LD motifs (positive set PS1). All the possible windows of length 10AA from the remaining regions of the above-mentioned six proteins were selected as the core windows of the initial negative set (NS1). This produced a set of 4020 windows from six proteins that formed NS1. To consider the importance of surrounding regions of LD motifs, 20AA residues flanking regions on each side of the scanning window were analysed.

### Feature extraction from protein sequences

From the set of aligned 18 windows with their flanking sequences, position frequency matrix (PFM) was constructed. If the flanking region of scanning window is shorter than 20AA (at N-terminal and C-terminal region) then the positions are filled up by a gap (‘-’). PFM was then normalised to produce Position Weight Matrix (PWM) using normalisation technique analogous to^73^. We only consider twenty IUPAC unambiguous AA codes (http://www.bioinformatics.org/sms/iupac.html) and gap (‘-’) for building PWM. We built PWM from the scanning core window (PWM_CoreSeq_) which consists of 10 residues, the two flanking regions each with 20 residues produces two other PWMs (PWM_UpSeq_, PWM_DownSeq_) and the whole segment (upstream flanking region + core window + downstream flanking region) of 50 (20 + 10+ 20) AA residues produces the additional PWM. Then, during the scanning of protein sequences, we matched the four PWMs with corresponding window segments to get the respective four matching scores^73^. We also considered the average values of the mapping score from the PWM of core window (PWM_CoreSeq_) and PWM of flanking regions (PWM_UpSeq_, PWM_DownSeq_). Thus, we generated five features for each window. While generating the scores from the core PWM (PWM_CoreSeq_) we used our previous knowledge of the properties of *bona fide* LD motifs ^19,31^. If there are no acidic residues (Asp or Glu) either at position 0 or 6, we assign the score zero to PWM_CoreSeq_. Proline has a tendency to break the helix. Consequently, if there were two consecutive prolines in core motif we also assigned 0 to PWMCoreSeq.

### Feature extraction from secondary structure (SS)

We predicted the secondary structure (SS) of the whole protein using PSIPRED^33^ against the NR database. Each residue in the 50AA window (core + flanking regions) was tagged as belonging to helix (‘H’) or coil (‘C’) or strand (‘E’). Gap (‘-’) was also considered for the windows near N/C-terminal of proteins. From the set of 18 windows that correspond to known LD motifs (with flanking regions), we constructed PFM matrices (analogously as mentioned in the previous section) based on SS annotation of residues. PFM was then normalized to PWM. We built PWM from the scanning core window (PWM_CoreSS_), the two flanking regions each with 20 residues produces two other PWMs (PWM_UpSS_, PWM_DownSS_) and the whole segment (upstream flanking region + core window + downstream flanking region) of 50 (20 + 10 + 20) AA residues produces the additional PWM. Using PWMs, we were able to generate five features from SS information in the analogous manner as explained in the previous section. In these cases, if the core motif part does not have any helical prediction, we assign zero to the core motif score from PWM_CoreSS_.

### Feature extraction using AAindex

From Amino Acid Index (AAindex) database^74^ three physiochemical properties were extracted: hydrophobicity^75^, volume, and electric charge^76^. For each of the 10 residues in a core window, we calculate the AAindex values of the above-mentioned three properties that produced 30 (3*10) features.

### Model Development

We generated an initial model based on the initial training data. Since this model is based on data derived from only six proteins and contains a very small number (18) of known LD motifs, we extended the training set by hypothetical LD motifs and additional negative data. For this, we used a procedure (explained below that, among other things, utilizes the initial model) that is likely to generate motifs highly similar to known LD motifs. Once the training set is expanded this way, we retrained the model as we used initially.

### The Initial Model

We extracted five features using primary sequence information, five features using SS information, and 30 features using AAindex for data-windows as discussed previously. Then we used a support vector machine (SVM) model ^77^ with linear kernel ^78^ to build a predictive model (M1). We used ‘svmtrain’ function of MATLAB 2012b with default parameter setting to build the model (there was no need to optimize parameters of the SVM model as the default setting provided an excellent performance).

### LD Motifs from Homologous Proteins

As we have very limited number of known LD motifs, we tried to increase that number using standard protein-protein BLAST (blastp) hits which are similar to motifs ^79^. We used the six proteins that contain the known LD motifs for the blastp program and selected the complete sequence of the proteins with the high score of BLAST hits (E-value:1e-7, bit score > 40). Then, we applied our M1 to identify the LD motifs from these proteins homologous to the six proteins that contained known LD motifs. In this way, we predicted 40 more LD motifs from these proteins. These additional 40 candidate LD motifs were also considered as correct and used for building our final model.

### Active Learning Dataset from PDB

We downloaded a culling set^28^ of proteins from the Protein Data Bank (PDB) to enhance our negative dataset-. We predicted SS of the full chain using PSIPRED. We built three independent models from the initial dataset based on five sequence features (M1_seq_), five SS features (M1_ss_) and 30 AAindex features (M1_aaindex_). For each of these models, we used an SVM model with linear kernel and default parameter setting.

We applied M1_seq_ to the culling set to predict windows with LD motifs. These windows formed the set S_seq_. Analogously, we generated sets S_ss_ and S_aaindex_ using M1_ss_ and M1_aaindex_, respectively. Our hypothesis was that a window that does not belong to the intersection of these three sets is less likely to contain LD motifs. So, we included such windows in the negative set. This has resulted in 2,279 additional negative data-windows used for building the final model.

### The Second Model (M2)

We extracted the features from all (18+40) positive and all (4020+2279) negative data-windows in the same fashion as discussed previously and we used an SVM with the linear kernel to build a predictive model (M2). We used ‘svmtrain’ function of MATLAB 2012b with default parameters setting to build the final model. This model predicts 13 new LD motif from human proteome. We applied a version of the 18-fold cross-validation (CV) to assess the model accuracy. We divided the negative set randomly into 18 disjoint subsets. At each step of CV, we excluded a different subset from the negative data and the window that corresponds to one of the 18 known LD motifs. Moreover, from the additional 40 positive data (windows) we excluded all windows from proteins homologous to the excluded one to which the known LD motif belongs. This last step is done in order to avoid dependent data in the training set.Then, the model is derived from the remaining data as described in the section above, and it was tested on the excluded data.

### The Final Model

We experimentally (*in vitro*) verified the 13 new LD motifs and found that four of them show a strong binding affinity (“Highly likely” category) towards their binding partners. So, we integrate these four motifs in the roster of true LD motif and build the final model following the same method described above. This final model predicts eight LD motifs. Three were new LD motifs and five were common to previously predicted 13 LD motifs by M2. Using CV approach, mentioned in the above section, the final model achieved over 88.88% sensitivity and accuracy of 99.97% (**Supplementary Table** 1).

### Validation of LDMF using Random Sets

To evaluate the robustness of our final model we tested it on random sequences generated by Sequence Manipulation Suite^80^. We generated 1,000 random sequences and applied the model to them. *LDMF* did not predict any LD motif in these sequences.

## Availability

*LDMF* is available at http://cbrc.kaust.edu.sa/ldmf/. For the result mentioned in this manuscript, we used the NR database for PSIPRED predictions. But for our online *LDMF* server, due to the prohibitive time required to obtain the results from the NR database, we used UNIPROT database for PSIPRED predictions.

### Proteins and Peptides

Human α-parvin-CH_C_ (residues 242-372), the FAT domain of human FAK (892-1052), and the rat GIT1 (647-770) were expressed as GST-fusion proteins in *E. coli* BL21 at using the expression vectors pGex 6P1, pGexP2, pGex-4T1, respectively. Bacteria were grown in LB medium. α-parvin-CH_C_ and FAT were expressed at 20°C overnight, whereas GIT1 was expressed for 6h at 30°C, α-parvin-CH_C_,FAT and GIT were purified as described previously ^21,22,81^.

Peptides were purchased from GenScript with and without FITC-Ahx N-terminal modification, with the following sequences: LD4 (SASSATRELDELMASLSD), LD2 (NLSELDRLLLELNAVQ), IBP2 (TPTQQELDQVLERISTMR), RGPA2 (GDDVLDKLLENIGHT), CH037 (AEDLDELLDEVESKFATPD), ICAL (DAALDDLIDTLGGP), FIP1 (SAGEVERLVSELSGGT), WHAMM (PGSMDEVLASLRHG), LPP (AEIDSLTSILADLESS), RGPA1 (EDVLDELLQYLGVT), CP071 (EAWDLDDILQSLQGQ), NCOA2 (SELDNLEEILDDLQNSQ), E41L5 (ATDELDALLASLTENLID), PCP2 (PTPEMDSLMDMLASTQ), NCOA3 (GDLDNLDAILGDLTSSD), RHGO7 (DIFPELDDILYHVKGMQ), P2R3A (SQEEIDKLLMDLESFSQ), CREB3 (SDWEVDDLLCSLLSPPA), CCDC158 (DPTRDLKQLLQELRSVIN), Scramble (LSDAMETSSLRDALE).

### Differential Scanning Fluorimetry

Experiments were performed in 20 mM HEPES pH 7.5, 150 mM NaCl, 2 mM EDTA, 1 mM TCEP. FAT, α-parvin-CH_C_ and GIT1 were used at a concentration of 10 μM. Protein stability was assessed for each peptide at 100 and 250 μM. SYPRO Orange was used as fluorescent dye at 1x the protein concentration. The samples were heated from 20°C to 95°C at a rate of 0.03°C/s on a LightCycler 480 II RT-PCR from Roche. When plotted as a function of the temperature, the fluorescence intensity resembles the classical sigmoidal function that is described by the Boltzmann equation. To estimate the melting temperature (Tm), a generalized sigmoid was fitted by least squares and the inflection point was computed.

### Direct anisotropy assay

Protein was serially diluted in buffer (20 mM HEPES pH 7.5, 150 mM NaCl, 2 mM EDTA, 2 mM DTT, 0.005% Tween-20) and labelled peptides were added at a final concentration of 0.1 μM. Fluorescence anisotropy was measured on a PHERAstar FS device (BMG Labtech) using a fluorescence polarization module 485/520/520, at room temperature. Fluorescence anisotropy was determined as: 1000*(I_//_ – I_+_)/(I_//_+2*I_+_), where I_//_ and I_+_ are parallel and perpendicular components of fluorescence intensity excited by parallel polarized light. Data were analysed with Origin software using a logistic fit.

### Anisotropy competition assay

First, FAT and α-parvin were titrated, and FITC-Ahx-labelled LD4 was added as described for the direct anisotropy assay. Competition for the LD4 binding site of FAT and α-parvin was then assessed as follows: the proteins were kept at a concentration corresponding to the K_D_ of their interaction with labelled LD4 (10 μM for FAT and 25 μM for α-parvin), in the presence of 0.1 μM labelled LD4. To that, each non-labelled peptide was added at 100 and 250 μM. When competing for the binding site, the unlabelled peptide displaces labelled LD4 resulting in a lower anisotropy. All measurements were performed as for direct anisotropy assay. Values are represented as a ratio to the point estimated to be the *K*d of the protein with LD4 labelled.

### Isothermal Titration Calorimetry

Proteins were dialysed in ITC buffer (20mM HEPES pH 7.5, 150mM NaCl, 1mM EDTA, 1mM TCEP). 1.5 ml of protein solution was placed in the cell at a concentration varying depending on the interaction from 50 to 150 μM for FAT and 125 μM for GIT1. Peptides were dissolved into the dialysis buffer to a concentration of 1 mM (for FAT interactions) or 1 to 1.25 mM (for GIT1) and placed in the injection syringe. Titrations were performed at 25 °C with an initial injection of 2 μl, followed by 23 injections of 10 μl. As a control, the peptide was titrated into the buffer and the resulting heats subtracted from the protein-binding curve. ITC was performed on a Nano ITC from TA Instruments, and data were fitted using NanoAnalyze Software. (For GIT1 to LD2 interaction, MicroCal iTC 200 from GE was used and data were fitted using Origin Software.

### Microscale Thermophoresis

Serial dilutions of proteins were prepared starting from 630 μM (GIT1), 560 μM (FAT) and 530 μM (α-parvin) concentrations, in reaction buffer (20 mM HEPES pH 7.5, 150 mM NaCl, 2 mM EDTA, 2 mM DTT, 0.05% Tween-20). Labelled peptides were added to a final concentration of 0.1 μM. The experiment was performed at 20 % LED power and 20, 40 and 60 % MST power in standard capillaries (GIT1) and MST Premium Capillaries (FAT and α-parvin) on a Monolith NT.115 device at 25 °C (NanoTemper Technologies). Thermophoresis and temperature jump were fitted using the K_D_ formula derived from the law of mass action on the provided NT analysis software.

### Nuclear Magnetic Resonance

Cells were grown with^15^N-labelled ammonium chloride dissolved in M9 minimal media solution, induced at OD=0.8 with 300uM IPTG and harvested after incubation overnight at 22⁏C. Protein samples were purified and NMR samples were prepared by dissolving the ^15^N-labelled protein in a 10% D_2_O/90% H_2_O solution with a monomer concentration of 100 *μ*L in a total volume of 500 *μ*L and pH of 7.5. LD motif-containing proteins were dissolved with FAT gel filtration buffer (20mM HEPES pH=7.5,150mM NaCl, 2mM EDTA and 2mM DTT). 2 *μ*L of 25 mM 2,2-dimethyl-2-silapentane-5-sulfonate (DSS) sodium salt was added as an internal chemical shift reference for ^1^H at 0 ppm. The samples were stable over the course of the NMR experiments. The ^1^H-^15^N HSQC titration experiments were performed at a temperature of 25° C using a Bruker Avance III 950 MHz NMR spectrometer equipped with a triple resonance inverse TCI CryoProbe. Spectra were acquired with 2048 (^1^H) × 200-256 (^15^N) complex points, a spectral width of 16 ppm for ^1^H and 40 ppm for ^15^N, and averaged for 36-88 scans depending on sample concentration. Changes in chemical shifts for ^1^H and ^15^N were measured in ppm (δ*H*and δN) where we multiplied the ^15^N shift changes by a scaling factor ⁀=0.14, and then calculated the summed Euclidean distance moved following this equation: d=|*δH*|+*α*|δN|^82^

### Data-driven Molecular Docking

The data-driven HADDOCK 2.1 protocol ^83^ was used to generate the models of complexes for FAT:CCDC158 and FAT:LPP. Crystal structures of FAT (1ow8 and 1ow7) were used for the modelling. Initial models for CCDC158 and LPP were modelled in helical form based on the LD4 peptide. The NMR chemical shift perturbation (CSP) data was used to define the residues, which could be potentially involved in the binding known as active residues. The residues 932, 934, 937, 1025, 1028, 1031, 1035 and 1040 were marked on FAT as active residues for FAT:CCDC158 and 956, 959, 962, 963, 964, 966, 991, 995 and 996 were marked on FAT as active residues for FAT:LPP. The CSP data was only used to define the binding site and not the binding poses. A total of 500 models were generated in the rigid-body docking step (it0), from which the top 100 according to the HADDOCK score were further refined with a simulated annealing interface (it1) and finally with a solvation step making a water-layer around the complex. Structures that were listed in the output clusters with best scores were further analysed using PyMol.

